# Clinical diagnosis of TIA or stroke and prognosis in patients with neurological symptoms: a rapid access clinic cohort

**DOI:** 10.1101/507012

**Authors:** Catriona Graham, David Bailey, Simon Hart, Aidan Hutchison, Peter Sandercock, Fergus Doubal, Cathie Sudlow, Andrew Farrall, Joanna Wardlaw, Martin Dennis, William Whiteley

## Abstract

**Background:** The long-term risk of stroke or MI in patients with minor neurological symptoms who are not clinically diagnosed with transient ischaemic attack (TIA) or minor stroke is uncertain.

**Methods:** We used data from a rapid access clinic for patients with suspected TIA or minor stroke and follow-up from four overlapping data sources for a diagnosis of ischaemic or haemorrhagic stroke, myocardial infarction, major haemorrhage and death. We identified patients with and without a clinical diagnosis of TIA or minor stroke. We estimated hazard ratios of stroke, MI and death in early and late time periods.

**Results:** 5,997 patients were seen from 2004–2013, who were diagnosed TIA or minor stroke (n=3604, 60%) or with other diagnoses (n=2392, 40%). By 5 years the proportion of patients who had a subsequent ischaemic stroke or MI, in patients with a clinical diagnosis of minor stroke or TIA was 19% [95% confidence interval (CI): 17–20%], and in patients with other diagnoses was 10% (95%CI: 8–15%). Patients with clinical diagnosis of TIA or minor stroke had three times the hazard of stroke or MI compared to patients with other diagnoses (HR 2.83 95%CI:2.13–3.76, adjusted age and sex) by 90 days post-event; however from 90 days to end of follow up, this difference was attenuated (HR 1.52, 95%CI:1.25–1.86). Older patients and those who had a history of vascular disease had a high risk of stroke or MI, whether or not they were diagnosed with minor stroke or TIA.

**Conclusions:** Careful attention to vascular risk factors in patients presenting with transient or minor neurological symptoms not thought to be due to stroke or TIA is justified, particularly those who are older or have a history of vascular disease.

## Introduction

National guidelines recommend rapid clinical assessment for patients with suspected stroke or transient ischaemic attack (TIA) in hospital-based clinics or emergency departments^1,2^ in order to prevent vascular events, chiefly stroke. Therefore, much depends on the clinical diagnosis of stroke or TIA. Risk prediction tools may aid stratification in patients with a diagnosis of TIA, but they may be less useful in all patients presenting with focal neurological symptoms due to a mixture of causes.^3^

The degree to which a bedside clinical diagnosis of stroke or TIA groups in clinical practice identifies a patient at higher subsequent stroke or myocardial infarction (MI) is unclear. Because doctors frequently disagree about the diagnosis of minor stroke or TIA in individual patients,^4–7^ in routine practice patients who are clinically diagnosed as having a low or medium risk of TIA may not be as extensively investigated, or offered secondary preventative treatments. In many areas of the UK and in other countries, stroke services do not have rapid access to advanced brain imaging; for patients in these regions the clinical diagnosis is the key step for patients in order to access secondary preventative medications and other strategies.

Although previous studies have examined the risk of recurrent vascular events in different groups of patients already diagnosed with TIA stratified with clinical risk models,^8^ or advanced brain imaging,^9^ few studies have examined whether the bedside clinical diagnosis of TIA or minor stroke discriminates between patients at a high and low risk of stroke or MI in all patients presenting to stroke services with neurological symptoms. We therefore sought to determine the effectiveness of the bedside clinical diagnosis of minor stroke or TIA, prior to imaging or other tests, for risk stratification in routine clinical practice in different groups of patients presenting with transient or minor neurological symptoms.

## Methods

### Data recording

We used anonymised data from a bespoke electronic health record (EHR) that was introduced in 2004 to a one-stop clinic for rapid assessment of patients with suspected TIA or minor stroke. Patients from across Lothian, a region of Scotland with a population of 810,000, were referred to the clinic by their general practitioner or other referrers (e.g. emergency department, eye hospital) either by letter or fax (2004–2007) or telephone or email (2007–present), and assessed face-to-face by a consultant or senior trainee in geriatrics or neurology who could order same day brain imaging [magnetic resonance imaging (MRI) or computerised tomography (CT)], carotid Doppler ultrasound or electrocardiography (ECG) where appropriate.^10^ Demographic and clinical details were recorded in the EHR, and imaging was ordered and recorded in the hospital radiology system. Because the EHR was used to produce immediate clinic letters, the majority of fields were completed in mutually exclusive categories during the clinic, and before the results of imaging were known.

### Population of interest

We included all patients at their first assessment in clinic whose primary residence was in Scotland. Each patient had a Community Health Index (CHI) number, an identifier that is unique to each individual in Scotland, which facilitates data linkage across multiple healthcare datasets.

### Diagnosis

Doctors recorded their diagnosis for each patient at the end of their clinical assessment before imaging and other tests, and did not change this field in the EHR in the light of these results. At the time of assessment, the doctor recorded a prior history of atrial fibrillation, cardiac, cerebral or peripheral vascular diseases, blood pressure, diabetes and the chief symptom of the presenting complaint. Diagnoses were coded as stroke, TIA, retinal artery occlusion, amaurosis fugax or other diagnosis. For each cerebrovascular diagnosis, the diagnostic certainty was recorded as *definite*, where no other diagnosis was contemplated, *probable* where a cerebrovascular diagnosis was the most likely, but other diagnoses were considered, *possible*, where a non-cerebrovascular diagnosis was most likely, though cerebrovascular diagnoses were considered, and *definitely not* when a non-cerebrovascular diagnosis was made. Our primary analysis compared those with a diagnosis of minor stroke or TIA, (i.e. definite or probable stroke, TIA or transient or permanent monocular blindness) versus those with other diagnoses (i.e. a non-cerebrovascular diagnosis, or those with a ‘possible’ diagnosis).

### Outcomes

We obtained follow-up data on subsequent ischaemic stroke, myocardial infarction, major haemorrhage or death from four overlapping data sources: (1) the TIA clinic EHR; (2) the Scottish Stroke Care Audit (SSCA) (strokeaudit.scot.nhs.uk), a nationwide rolling audit of all episodes of in- and out-patient stroke care; (3) the Scottish Morbidity Record (SMR01), a nationwide register of all hospital admission; and (4) the General Registry Office, a nationwide death register.

We defined subsequent ischaemic stroke as the first record of ischaemic stroke in: (1) a new record of definite or probable stroke in the TIA clinic EHR at any time after the index event; (2) ischaemic strokes in the Scottish Stroke Care Audit where the stroke pathology was recorded as ischaemic or due to haemorrhagic transformation of an infarct, and the date of onset of the stroke was >7 days after the date of onset of symptoms or assessment date (whichever was later) in the TIA clinic EHR; and (3) the Scottish Morbidity Record, where an admission or death was due to ischaemic stroke (i.e. ICD-10 codes I63, and ICD-9 codes - 434) and the admission was >7 days after the date of onset of symptoms or assessment date in the EHR. We used a delay of >7 days in nationally collected datasets as our primary definition of recurrent stroke, though explored delays of 0 and 30 days in sensitivity analyses. We defined subsequent myocardial infarction as admission to hospital recorded in the SMR01 or death with ICD-10 codes I21 and I22, or ICD-9 code −41 at any time. We defined subsequent major haemorrhage as the ICD-10 codes −43, −44, H43, I60, I61, I62, K250, K254, K260, K264, K266, K272, K290, and R040.

### Statistical analyses

We performed statistical analysis with an anonymised dataset. In the primary analysis patients were divided into two groups by the level of certainty of their baseline diagnosis of stroke or TIA (definite/probable versus possible/definitely not), which we refer to as ‘minor stroke or TIA’ versus ‘other diagnoses’. We followed up patients to the first record of ischaemic stroke or MI in the EHR, SSCA or SMR, or haemorrhage, or death. Patients were censored if they died from a cause other than stroke or MI. We plotted cumulative incidence curves, compared the curves with log-rank tests and used the survivor function to estimate the absolute risk of ischaemic stroke or MI at 5 years. We compared the two groups with Cox regression, after checking the proportional hazards assumption and reported the hazard ratios (HRs) and their 95% CI. We adjusted the HRs initially for age and sex, and further for history of stroke or TIA, MI, angina, peripheral vascular disease, diabetes, cardiac failure and AF. We looked for significant linear multiplicative interactions of diagnosis with age, sex, clinical experience of the examining clinician, AF, prior vascular diseases, the chief presenting complaint and delay from clinical event to assessment, to determine whether a clinical diagnosis was more or less useful in different types of patients.

## Results

Between December 2005 and May 2013, 37 different doctors saw 5,997 patients referred with an episode of neurological disturbance that was suspected to be due to a transient ischaemic attack or stroke. The patients’ characteristics are shown in Table 1. The median age at assessment was 68.5 years (IQR 58–77.5), 49.5% were women, and the median delay to assessment from symptom onset was 5 days. Patients were seen by neurologists (19%) or stroke physicians (81%), roughly half by a trainee (52%) and half by a consultant (48%). 3604 (60%) patients had a clinical diagnosis of stroke, TIA or monocular blindness, and 2392 (40%) had other diagnoses. The clinically definite or probable cerebrovascular diagnoses were: stroke (1554, 43%), TIA (1582, 44%), and transient or permanent monocular blindness (468, 12%).

**Table 1.** Characteristics of patients seen within the stroke/TIA clinic, 2004-2013

| Diagnostic certainty of stroke or TIA |  | p-value for difference |
| --- | --- | --- |
| Possible/not | Definite/probable |  |

|  | N | % | N | % |  |
| --- | --- | --- | --- | --- | --- |
| **Number of patients** | 2392 | 100 | 3604 | 100 |  |
| **Gender** |  |  |  |  |  |
| Female | 1284 | 54 | 1681 | 47 | <0.0001 |
| Male | 1108 | 46 | 1923 | 53 |  |
| **Age group** |  |  |  |  |  |
| <50 | 440 | 18 | 197 | 5 | <0.0001 |
| 50 – 70 | 1149 | 48 | 1475 | 41 |  |
| >70 | 803 | 34 | 1932 | 54 |  |
| **Speciality** |  |  |  |  |  |
| Missing | 83 | 3 | 106 | 3 |  |
| Neurology | 552 | 23 | 558 | 15 | <0.0001 |
| Stroke | 1757 | 73 | 2940 | 82 |  |
| **Seniority** |  |  |  |  |  |
| Missing | 83 | 3 | 106 | 3 |  |
| Consultant | 1088 | 45 | 1705 | 47 | 0.226 |
| Registrar | 1221 | 51 | 1793 | 50 |  |
| **Symptoms** |  |  |  |  |  |
| Speech disturbance | 274 | 11 | 888 | 26 | <0.0001 |
| Visual disturbance (1 eye) | 162 | 7 | 487 | 14 |  |
| Visual disturbance (2 eyes) | 288 | 12 | 238 | 7 |  |
| Motor | 416 | 17 | 1298 | 36 |  |
| Sensory | 452 | 19 | 381 | 11 |  |
| Dizziness | 341 | 14 | 189 | 5 |  |
| Headache | 41 | 2 | <10 | 0.2 |  |
| Lightheaded | 43 | 2 | <10 | 0.2 |  |
| Loss of consciousness | 45 | 2 | <10 | 0.1 |  |
| Other | 324 | 14 | 104 | 3 |  |
| None | 6 | 0.25 | 1 | - |  |
| **Prior stroke or TIA** |  |  |  |  |  |
| Missing | 10 | 0.4 | 8 | 0.2 |  |
| No | 1984 | 83 | 2779 | 77 | 0.052 |
| Yes | 398 | 17 | 817 | 23 |  |
| **Prior angina or MI** |  |  |  |  |  |
| Missing | 10 | 0.4 | 8 | 0.2 |  |
| No | 1972 | 82 | 2711 | 75 | <0.0001 |
| Yes | 410 | 17 | 885 | 25 |  |
| **Prior peripheral vascular disease** |  |  |  |  |  |
| Missing | 10 | 0.4 | 8 | 0.2 | 0.0003 |
| No | 2297 | 96 | 3395 | 94 |  |
| Yes | 85 | 4 | 201 | 6 |  |
| **Prior atrial fibrillation** |  |  |  |  |  |
| Missing | 10 | 0.4 | 8 | 0.2 |  |
| No | 2254 | 94 | 3244 | 90 | <0.0001 |
| Yes | 128 | 5 | 352 | 10 |  |
| **Prior diabetes** |  |  |  |  |  |
| Missing | 10 | 0.4 | 8 | 0.2 |  |
| No | 2143 | 90 | 3152 | 87 | 0.006 |
| Yes | 239 | 10 | 444 | 12 |  |

Patients with clinical diagnosis of minor stroke or TIA were more likely (all P<0.0001) than those with other diagnoses to be men (53% vs 46%), to be older (median age 72 yrs vs 64 yrs), or to have seen a specialist in stroke rather than neurology (82% vs 73%). Patients with a clinical diagnosis of minor stroke or TIA had a greater frequency (all p<0.001) of prior stroke or TIA (23% vs 17%), angina or MI (25% vs 17%), peripheral vascular disease (6% vs 4%), atrial fibrillation (8% vs 5%) and diabetes (12% vs 10%). There were statistically significant, though small, differences between the groups in mean systolic blood pressure at assessment (148 vs 144 mmHg, p<0.0001) and median time to assessment (5 vs 6 days, p<0.0001) though not diastolic blood pressure (82 vs 82mmHg P=0.210). (Table 1)

During follow up [median 4.2 yrs (interquartile range 2.4 – 5.8 yrs), longest follow-up 8.2 yrs], 655 patients had an ischaemic stroke, and 116 had a myocardial infarction. Stroke or MI occurred in 630 patients of the 3604 patients with a diagnosis of acute stroke or TIA, and 206 in the 2392 patients with other diagnoses. (Figure 1). In the whole group, MI and stroke respectively occurred in 34 and 306 patients from 0–90 day, and 306 and 349 patients 90 days-end of follow up. The hazards ratio for stroke or MI changed over the follow up period (p<0.05), and therefore we report them from 0–90 days, and 90 days–end of follow up.

**Figure 1.**
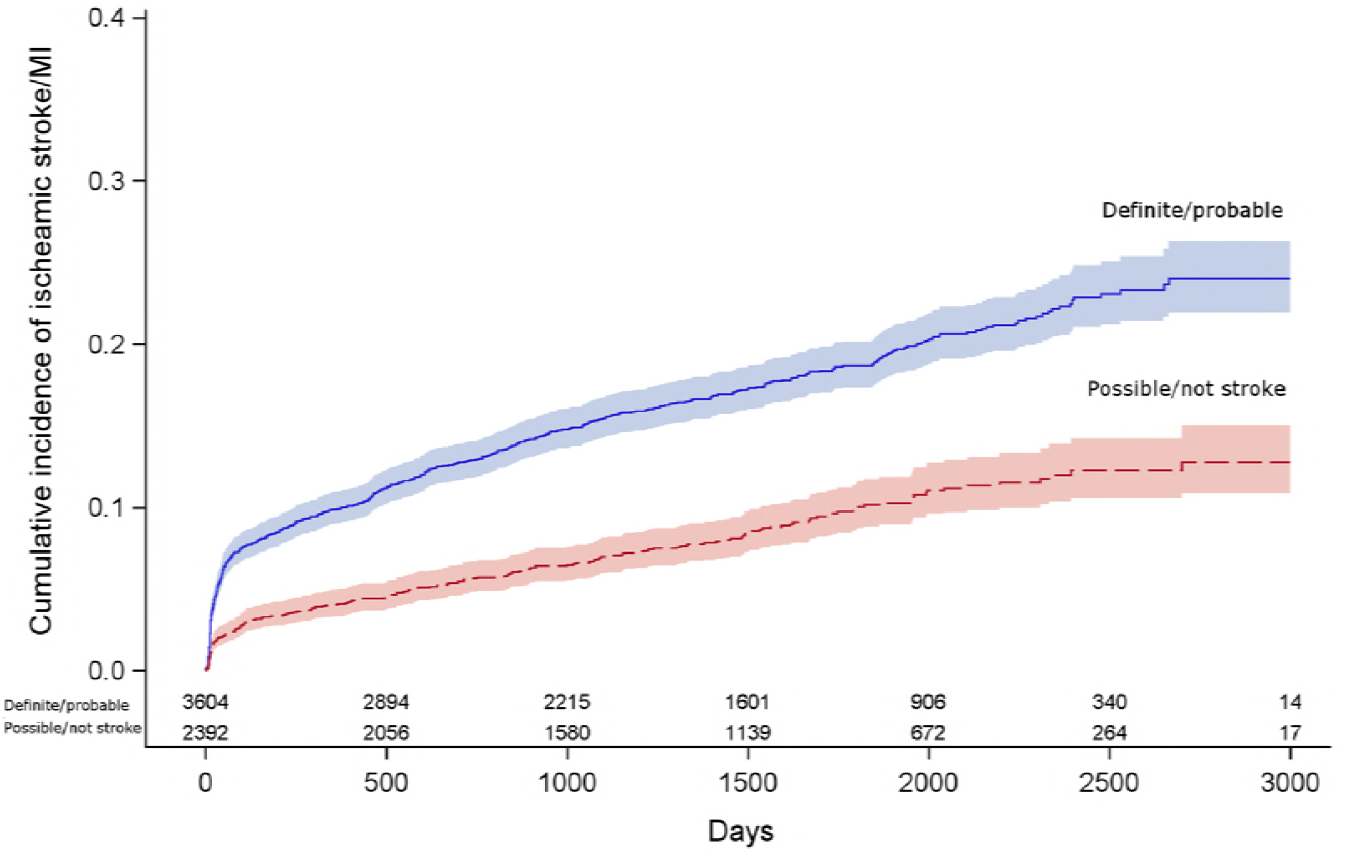
Cumulative incidence curves of ischaemic stroke or myocardial infarction in follow up in patients with a diagnosis of definite or probable TIA/stroke versus possible or not TIA/stroke, with 95% CI.

From 0–90 days from symptom onset (or where this was unclear, assessment in clinic), the hazards of stroke or MI were approximately 3 times higher in patients with stroke or TIA, compared with those with other diagnoses (263 vs 62, HR: 2.88, 95%CI 2.19–3.80), and attenuated slightly after adjustment (HR 2.76, 95% CI: 2.08–3.68). From 90 days to the end of follow up, this hazard ratio was less than in the early period (367 vs 144, 1.81, 95%CI: 1.49–2.20) and attenuated after adjustment (HR1.40, 95% CI: 1.14, 1.70). (Table 2)

**Table 2.** The unadjusted and adjusted hazard ratios for ischaemic stroke or MI, major haemorrhage or death during follow up comparing patients with a diagnosis of definite or probable stroke or TIA to patients with another diagnosis. As the hazards for recurrent stroke and MI were not proportional over time, we present these separately for the periods 0–90 days and 90 day to end of follow up.

|  | Recurrent ischaemic stroke or MI |  | Major haemorrhage | Death |
| --- | --- | --- | --- | --- |
|  | 0-90 days | 90 day – end of follow up | 0 days–end of follow up | 0 days–end of follow up |
| **Unadjusted HR** | 2.88 (2.19–3.80) | 1.81 (1.49–2.20) | 1.60 (1.15–2.21) | 1.54 (1.34–1.76) |
| **Adjusted<sup>1</sup> HR** | 2.83 (2.13–3.76) | 1.52 (1.25–1.86) | 1.20 (0.86–1.68) | 1.05 (0.92–1.20) |
| **Adjusted<sup>2</sup> HR** | 2.76 (2.08–3.68) | 1.40 (1.14–1.70) | 1.17 (0.83–1.63) | 0.98 (0.86–1.13) |

1. Adjusted for age and sex
2. Adjusted for age, sex, MI, stroke, TIA, atrial fibrillation, angina, CF, PVD and diabetes

There was no strong evidence that the increased hazard of stroke or MI up to 90 days in patients with minor stroke or TIA relative to patients with other diagnoses differed (i.e. no statistically significant interaction) by the speciality of the assessing clinician, a patient’s history of AF, previous occlusive vascular event, hypertension at assessment or by the chief presenting symptom. (Figure 2) However, there was interaction with age (<50 yrs, HR: 8.91, 95%CI: 3.12–23.9; >70 yrs 1.97, 1.32-2.95, p_interaction_=0.015), because stroke and MI risk in the youngest patients with other diagnoses by 90 days was very low (1%) in comparison to older patients.

**Figure 2.**
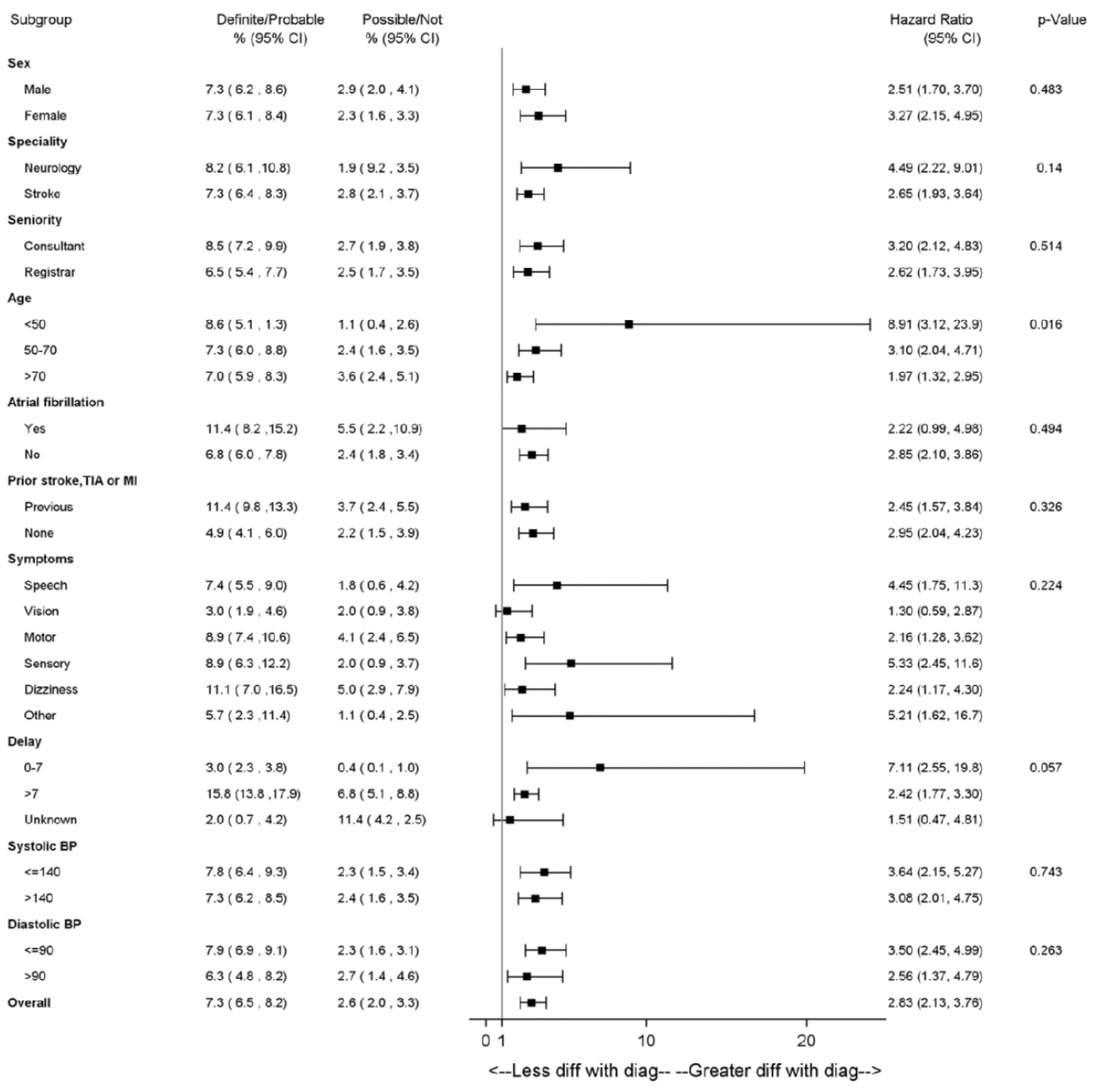
The risks of ischaemic stroke or MI by 90 days in patients with a clinical diagnosis of minor stroke or TIA versus those with other diagnoses in different groups of patients presenting with transient or minor neurological symptoms. Apart from the analyses of age and sex, all associations are adjusted for age and sexP-values indicate the significance of multiplicative interaction tests, i.e. the probability that differences in the HR between different groups of patients is due to chance. N=5997.

From 90 days to the end of follow up (Webfigure Figure 1) the increased hazards of MI or stroke in patients with minor stroke or TIA relative to those with other diagnoses was lower in patients with a history of prior vascular disease than those without [prior occlusive vascular disease HR:1.1 (0.85–1.43), no prior vascular disease 2.05 (1.52–2.79) p_interaction_=0.001]. This was because the risk of MI or ischaemic stroke in those with prior vascular disease was high whether or not the clinical diagnosis of the presenting event was acute stroke or TIA (194/1194,16.3% vs 85/616, 13.8%).

In sensitivity analyses, whether we considered a recurrent strokes at 0 days or greater after presentation (adjusted 0-90 day HR 3.36, 95%CI: 2.78–4.07), or at 30 days or greater after presentation (3.05,1.90–4.88) led to no great difference in the hazards of recurrent events. (webtable 1).

By 5 years, the absolute risk of stroke or MI in patients with a clinical diagnosis of stroke or TIA was 19% (95%CI: 17-20%), compared with 10% (95%CI: 9-12%) in patients with other diagnosis. Changing the timing of the definition of recurrent stroke made little difference to the absolute risk of recurrent stroke or MI by 5 years in patients with diagnoses other than stroke (0 days 13%, 95%CI: 12–15%; 30 days 9%, 95%CI: 7–10%), although reducing the time from presentation to recurrent stroke did increase the absolute risk of stroke or MI by 5 years in patients with a diagnosis of stroke or TIA [0 days 28%, 95%CI: 26–30%; 30 days 15%, 95%CI: 14–17%]. Further stratification of the diagnosis of clinical events into definite, probable, possible or other diagnoses led to further gradation of absolute risk (webfigure 2).

Over the course of follow up, 105 patients had a haemorrhagic stroke, 53 had an extracranial bleed, and 1002 patients died. Patients with a diagnosis of stroke or TIA had 60% greater hazard of major haemorrhage compared to patients with other diagnoses, which attenuated after adjustment [unadjusted HR 1.60, 95%CI:1.15–2.21; adjusted HR 1.17 95%CI: 0.83–1.63]. Patients with a diagnosis of stroke or TIA had approximately 50% greater hazard of death compared to patients with other diagnoses but this attenuated after adjustment [unadjusted HR 1.54, 95%CI:1.34–1.76; adjusted HR 0.98 95%CI: 0.86–1.13]. (Table 2)

## Discussion

There are patients presenting to TIA clinics who are at high risk of stroke or MI whether or not they were diagnosed with stroke or TIA: older patients, and those with a history of vascular disease. Therefore a strategy that might be effective in older patients, or patients with a history of stroke or MI who develop minor neurological symptoms, would be to use the presentation with symptoms to optimise preventative therapies whatever the clinical diagnosis.

Patients presenting with minor or transient neurological symptoms who were not diagnosed with TIA or minor stroke had a moderate risk of subsequent stroke or MI. Some of these patients had diagnoses such as migraine, syncope and more rarely epilepsy or brain tumours),^11–13^ though the majority had symptoms that were not easily explained by a clear alternative diagnosis (‘transient neurological attacks’, TNA). In previous studies, the relative risk for all occlusive vascular events (MI or ischaemic stroke) in the long term were similar in patients with TIA versus non-TIA in clinical (OR 1.2, 95%CI: 1.05–1.41) and some research settings (HR 1.0, 95% CI:0.8–1.2).^14–16^ The one year absolute risk of all occlusive vascular events in patients with symptoms that were not clear enough to make a diagnosis of TIA in the SOS-TIA study was similar to those with a definite diagnosis of TIA but normal brain imaging (2.18%, 95%CI 0.71–6.66 vs 2.78%, 1.65–4.65), although less than in those with a TIA and abnormal imaging (5.74%, 2.62–12.34).^17^ Therefore a key challenge for practice and research will be to determine whether a strategy of enhanced risk stratification in all patients with transient neurological attack (for example brain MRI with diffusion weighted imaging) is superior to current practice, or to more aggressive management of vascular risk in all patients with transient neurological attacks.

There are limitations to these conclusions.

We measured recurrent stroke or myocardial infarction recorded in electronic health records, therefore we had no opportunity to review the primary record of each case of recurrent stroke or MI. Although we are likely to have under-ascertained stroke (chiefly strokes that presented only to primary care or in patients who emigrated), the absolute risks of stroke in patients with a diagnosis of acute stroke or TIA were approximately the same as in studies with face-to-face or telephone follow up. We used an interval of 7 days to define recurrent stroke, because we were uncertain whether our different electronic sources had sufficient temporal resolution to differentiate different stroke events within one week; shorter and longer intervals made little difference in the relative hazards of stroke in patients with different level of diagnostic certainty, though a shorter interval did lead to a greater estimate of the absolute risk. The accuracy of Scottish records has been recently reviewed for MI and stroke and found to be reasonable. ^18,19,20^ Therefore, electronic health records are a suitable means to provide long-term follow up of large clinical cohorts, and provide plausible estimates of risk.

We use clinical bedside diagnosis, which will inevitably be influenced by factors of training (for example neurologists were more likely to make non-cerebrovascular diagnoses), or prior medical history. Because we tested our hypotheses in a single centre, there may be less variation in the diagnosis of stroke between the doctors in our centre than there would be between doctors across many centres with different practices. However, we tested diagnosis from a large number of clinicians at different levels of training; our practices are similar to others in the UK, and therefore we believe results are generalisable to those with a similar practice.

Our study has a number of advantages. The study included a large number of patients, and therefore small biases in misclassification would not have led to any important differences in our conclusions. We followed up patients for very long periods, with little loss to follow up as we recorded all events in Scotland. We used diagnoses made in clinical practice, rather than in research studies, therefore our results have relevance to practice in the UK and similar countries. Whilst the delay from the onset of symptoms to assessment in our clinic was 5 days, the delay from referral to clinic assessment is about 3 days^10^ Our clinic is representative of practice in Scotland: 94% of patients with TIA or minor stroke are seen in a clinic within 4 days of referral, compared with 83% across the whole country. (strokeaudit.scot.nhs.uk) Therefore our estimates of the proportions of patients with events are generalizable to services with similar practice.

## Conclusion

Although it is well known that patients with a diagnosis of TIA or minor stroke are at a high risk of recurrent stroke or MI, older patients, and patients with a history or vascular events who are not diagnosed with a TIA in rapid access TIA clinics also have a moderate to high risk of stroke in the long term. Careful attention to the control of vascular risk in these patients is justified.

## Acknowledgements

We would like to acknowledge the Scottish Stroke Care Audit

## Contributions

CG and WW designed and performed the analyses, DB linked nationally acquired datsets, SH, FD, PS, CS, AF, JW, MD and WW acquired patient’s data, AH contributed to informatics. WW drafted the paper, and is the guarantor. All authors revised the paper for important intellectual content, gave approval for the final version of the manuscript.

